# The evolution of embodied postural dynamics underlies behavioral diversity in benthic sessile chordate

**DOI:** 10.1101/2025.08.28.672800

**Authors:** Oleg Tolstenkov, Sissel Norland, Marios Chatzigeorgiou

**Author notes:** Contributed equally.

## Abstract

Benthic animals live on or in the seafloor, a habitat known as the benthic zone. These bottom-dwelling organisms account for the largest fraction of animal diversity in our seas, playing vital roles in nutrient cycling, sediment stabilization and the broader food webs. Historically, the behavioral repertoire of benthic animals attached permanently to the sea floor has been considered as extremely limited (if at all present), reflecting an evolutionary adaptation to their sessile lifestyle and simplified nervous systems. Here, we overturn this view by studying the spontaneous and stimulus evoked behavioral repertoire of the basal chordate *Ciona intestinalis* which has a benthic sessile adult stage. We found that *Ciona* adults transition between three main postural engagement behavioral states. These are defined by different body part kinematics and postural dynamics characterised by changes in body shape configuration as opposed to locomotion. Transitions between states can occur probabilistically in spontaneously behaving animals suggesting that they can be internally generated by the simple brain of an adult *Ciona*. Importantly, using spatiotemporal embedding of postural features, we constructed a behavioral space for *Ciona* adults, which was divided into 18 stereotyped behavioral modes revealing the unappreciated richness of adult *Ciona* behaviors. Our analysis showed that *Ciona* adults can deploy in a selective manner these behavioral modules to elicit distinct responses to different mechanical stimuli, suggesting the presence of a sensory context dependent regulation of behavior in these benthic organisms. Our study raises the possibility that embodied postures is underlie the evolution of behavioral repertoires in sessile benthic animals.

## INTRODUCTION

The fossil record suggests that a significant fraction of the Ediacaran bioata (575-542 Mya) was composed of benthic marine organisms with a tubular, oval or fond-like shaped form(*1–3*). A lot of these were sessile, anchored to the benthos (*4*). They would acquire their food either via osmotrophy or suspension feeding(*2, 5*) and some would show a very limited behavioral repertoire (e.g. aligning with the current flow)(*6*).

The Ediacaran biota abruptly disappeared 542 Mya, likely because of mass extinction and/or through biological interactions with the animals of the Cambrian explosion. Rapidly evolving animals in the Cambrian explosion included epibenthic sessile species capable of predation or filter-feeding such as stem-group ctenophores, cnidarians, sponges, brachiopods and tunicates(*7–14*).

Numerous sessile benthic organisms exist amongst extant metazoans. The majority arise from a pelagobenthic cycle, where a planktonic larva undergoes the transformative process of metamorphosis to become a sexually mature sessile, epibenthic adult. A biphasic lifecycle is often associated with dramatic morphological, physiological and behavioral changes between larval and adult stages(*15, 16*). These changes can be beneficial in several ways including occupying distinct ecological niches and specializing in different activities. For example, the freely swimming larvae can take advantage of marine currents to travel far away from the site of release(*17*), while the adults can be benthic, sessile filter feeders, which grow and spawn a large number of eggs and sperm. Larval behaviors and neurophysiology are being extensively studied across a broad diversity of marine species with a biphasic life cycle(*18–29*). However, we have a much more fragmentary understanding of the adult behavioral repertoire and sensory physiology, since studies of the benthic (and often sessile) adult stage are sparse(*30, 31*).

Ascidians are marine invertebrates with a biphasic life cycle. As the sister group to vertebrates, they have become important models for studying chordate development, physiology, behavior and ecology(*32*). Solitary ascidians like the emerging model organism *Ciona intestinalis*, have a freely swimming lecithotrophic larva, which eventually settles and metamorphoses to a benthic, sessile and filter feeding adult. The central nervous system of *Ciona* larvae share key characteristics with other chordates and thus it is the focus of intense research across scales from molecules to behaviors (*18, 19, 29, 33–44*). During the process of metamorphosis, most larval structures degenerate, and they are replaced by adult tissues(*45–47*). These adult structures originate primarily from various pools of undifferentiated progenitor cells (rudiments) present in the larvae(*47–51*). Despite being an order of magnitude larger in terms of number of neurons than the larval nervous system, the adult brain of *Ciona* is considered a relatively uninteresting organ, potentially due to a misconception that a sessile organism does not need a brain since it is not motile. Adult ascidians brains are adapted to sessility, evolving sensory organs and reflexes suitable to their benthic environmental niche (*42, 52–56*). The nervous system of the adult *Ciona* is composed of the cerebral ganglion and the neural gland, which make up the CNS and the dorsal strand plexus, the visceral nerves and a more diffuse network of anterior and posterior nerves that make up the peripheral nervous system(*55–59*).

To date, a small number of neurophysiological studies have been performed in adult ascidians. Using primarily electrophysiological assays they have shown that ascidian adults can detect and respond to stimuli like light, water flow, acute mechanical poke, vibrations of different frequencies and they can distinguish between particles of different sizes that pass by their siphons (*55, 56*). However, this physiological data is missing behavioral correlates. Historically, animal behavior tracking tools required manual annotation of numerous video frames and early tracking software focused either on following the center of mass of freely moving animals without providing postural information or extracting the midline and/or contour of a few predefined body shapes (*60–62*). The advent of convolutional neural networks and markerless pose tools for tracking a huge diversity of animals, such as DeepLabCut(*63, 64*), SLEAP(*65*) and others(*66, 67*) offers an unprecedented opportunity for the automatic extraction of ascidian adult body’s outline capturing most of the pose information collected in video recordings.

Here we leverage time-lapse video recordings of *Ciona* adults in combination with the markerless pose estimation tool DeepLabCut(*63*) to map the *Ciona intestinalis* adult behavioral repertoire. We obtained a detailed overview of shape-based features and body part kinematics. Using these shape features and body part kinematics we find that *Ciona* adults spend most of their time in one of three major behavioral states. These states differ from each other in terms of postural features and kinematics, duration and frequency of occurrence. Using dimensionality reduction, we derived lower dimensional representations of body postures, which we define as “Eigencionas” (ECs). With these ECs, we can explain the majority of postural variance in the *Ciona* adults. In addition, we exposed the animals to two types of mechanical stimuli, which we found elicited distinct sensory responses as quantified by postural features/kinematics, ECs and spatiotemporal embedding.

## RESULTS

### Parametrization of adult *Ciona* using interpretable features reveals three behavioral states

Using an inexpensive camera setup, we recorded high-resolution movie data of adult *Ciona intestinalis* individuals performing spontaneous and stimuli evoked behaviors. To automatically extract *Ciona* adult posture, we trained a DeepLabCut model, where we defined multiple points (18 points) along the animal’s body (Figure 1A; Movie S1). By reconstructing the outline of the *Ciona* adult body, we could follow the opening and closing of the atrial and oral siphons, as well as changes in the posture of the trunk shape. Outline coordinates were converted to kinematic and postural metrics for the different animal body regions (Figure 1B). In addition, we generated 30 Elliptical Fourier Descriptors (EFDs) to efficiently capture both gross and subtle changes in body posture over time (Figure S1A -S1C).

**Figure 1.**
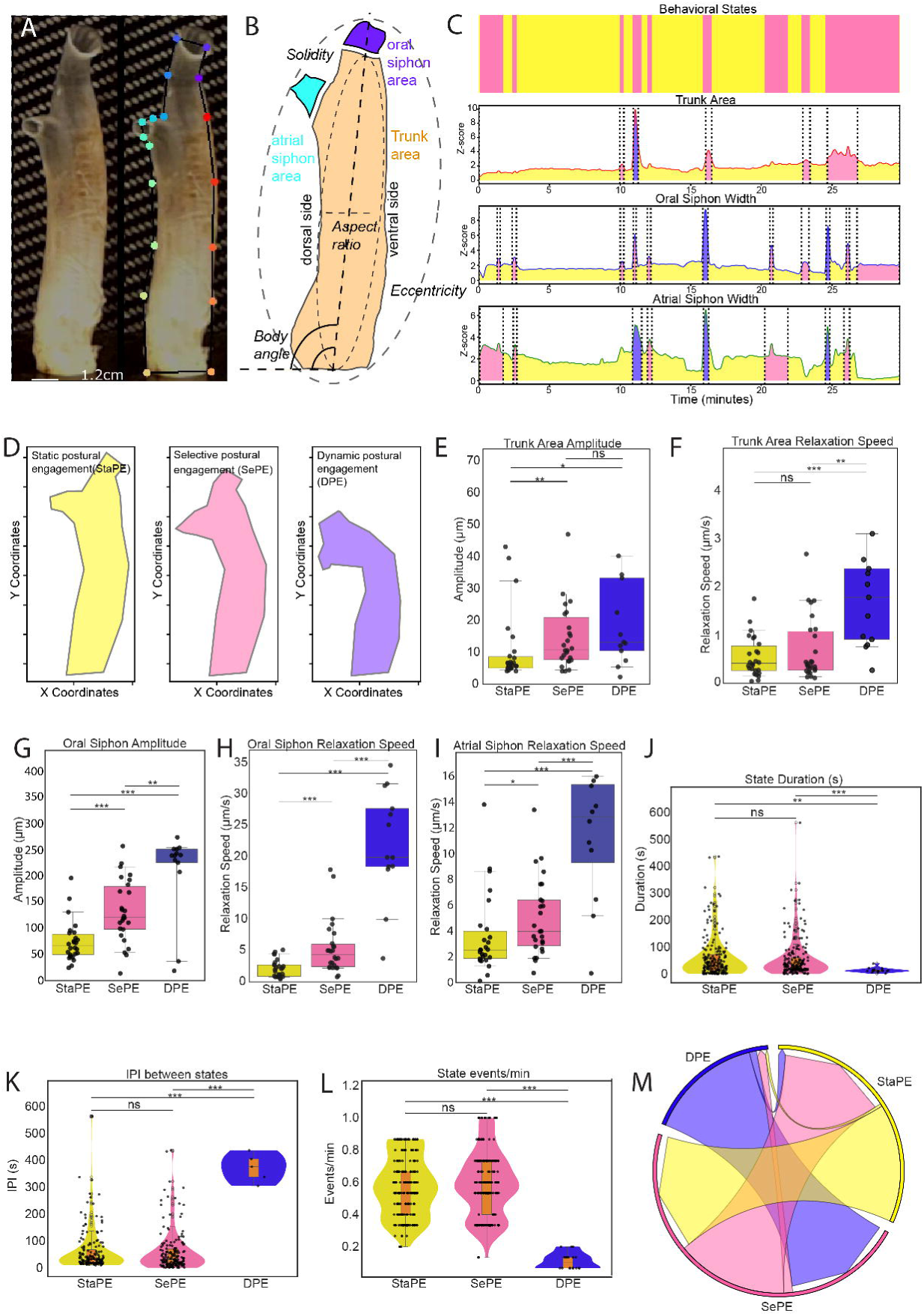
Parametrization of adult *Ciona* using interpretable features reveals three behavioral states. (A) Video frame of a *Ciona intestinalis* adult without (left) and with (right) DeepLabCut based annotation of key contour landmarks (see also Movie S1). (B) Schematic of adult *Ciona* body parts and morphological features used in our analysis. (C) Ethogram (top panel) of a single *Ciona* adult video-recorded for 30 minutes exhibiting spontaneous behaviours. Coloring corresponds to the behavioral modules defined based on threshold values of trunk area, oral siphon width and atrial siphon width. The corresponding z-scored time series of these features for this individual *Ciona* are shown in the lower three panels. (D) Reconstruction of *Ciona* contours from individual video frames that have been classified as belonging to one of the three states: Dynamic postural engagement (DPE), Selective postural engagement (SePE), Static postural engagement (StaPE). (E-I) Box plots quantifying different postural features and body part kinematics used in the characterization of the three behavioral states. Mann Whitney U tests were used for statistical analysis (ns p>0.05; * p<0.05; ** p<0.005; *** p<0.0001). (J-L) Violin plots quantifying the temporal features in each of the three behavioral states. (M) Chord diagram showing the transitions between the three behavioral states in spontaneously behaving animals.

Spontaneously behaving *Ciona* adults show variability in behavior across time (e.g. Movie S1). In other animals, this variability is generated through rapid shifts in behavior as animals transition between different behavioral states (*68, 69*). A plethora of behavioral states have been documented in the literature for animals that can freely locomote, however a lot of these states (e.g. roaming, dwelling, foraging, escape) are not applicable to sessile organisms like the benthic *Ciona* adult which are fixed to one place permanently. To behave, sessile animals do not rely on locomotion but rather they engage their body’s postural control system aimed at actively maintaining specific postures and to transition between them. We define this process as postural engagement.

Analyzing the spontaneous behavior of adult *Ciona* we identified three behavioral states, which we classified as Dynamic postural engagement (DPE), Selective postural engagement (SePE), Static postural engagement (StaPE) (Figure 1C, 1D; Figure S1D). These states were defined by the synchronization of the contraction events in the three body regions characterized by the difference in various postural and kinematic parameters of the three body regions (Figure 1E-I; Figure S1E-P). In the case of the trunk, area amplitude and relaxation speed best differentiated the states (Figure 1E and 1F; Figure S1E). For the oral siphon region aperture amplitude and relaxation speed were the most informative features (Figure 1G and 1H; Figure S1F-I). Finally, for the atrial siphon, relaxation speed was the most informative feature (Figure 1I; Figure S1J-N). On occasion, we noticed that the adult *Ciona* bodies acquired a slightly bent posture but neither body angle nor dorsal/ventral ratio metrics showed a significant difference (Figure S1O and S1P) between states during spontaneous behavior. Notably, a Hidden Markov Model approach segmented behavioral time series data in a manner that was generally consistent with our manual definition of states (Figure S1Q).

We next characterised the temporal features of the behavioral states. We found that the StaPE state had the shortest duration (Figure 1J) and the lowest frequency of occurrence amongst the three states (Figure 1K and 1L). The temporal dynamics of the DPE and SePE states were not significantly different from each other (Figure 1J-1L). We also analysed the transition probabilities between states. Animals in the intermediate SePE state could transition to either DPE or StaPE. However, we failed to identify direct transitions between the DPE and StaPE states. This is indicative of a stereotyped sequence of transitions in behavioral states in the absence of an external sensory stimulus (Figure 1M).

### Mechanical stimuli elicit distinct behavioral responses

*Ciona* adults are equipped with mechanoreceptors on their siphons and these are sensitive to water flow and poking as demonstrated by electrophysiological studies(*55, 56*). Here, we recorded and analyzed the behavioral responses of *Ciona* adults in response to mechanical poking and water flow (Figure 2A and 2B; Figure S2 A-D; Movies S2 and S3). We found that a mechanical poke outside the oral siphon elicited a strong defensive response by entering a DPE state and performing a full contraction (Figure 2A and 2C blue bars). A DPE state was observed with significantly higher probability immediately following a poke stimulus compared to water flow stimulated and control unstimulated animals (Figure 2D). When mechanically poked animals exited the DPE state, they showed a higher probability of entering a StaPE (Figure 2C (pink bars) and 2D). This is in sharp contrast to what we observed in controls (no-stimulus) where this transition was never observed. This suggests that acute mechanical stimulation can introduce plasticity in the state transition dynamics.

**Figure 2.**
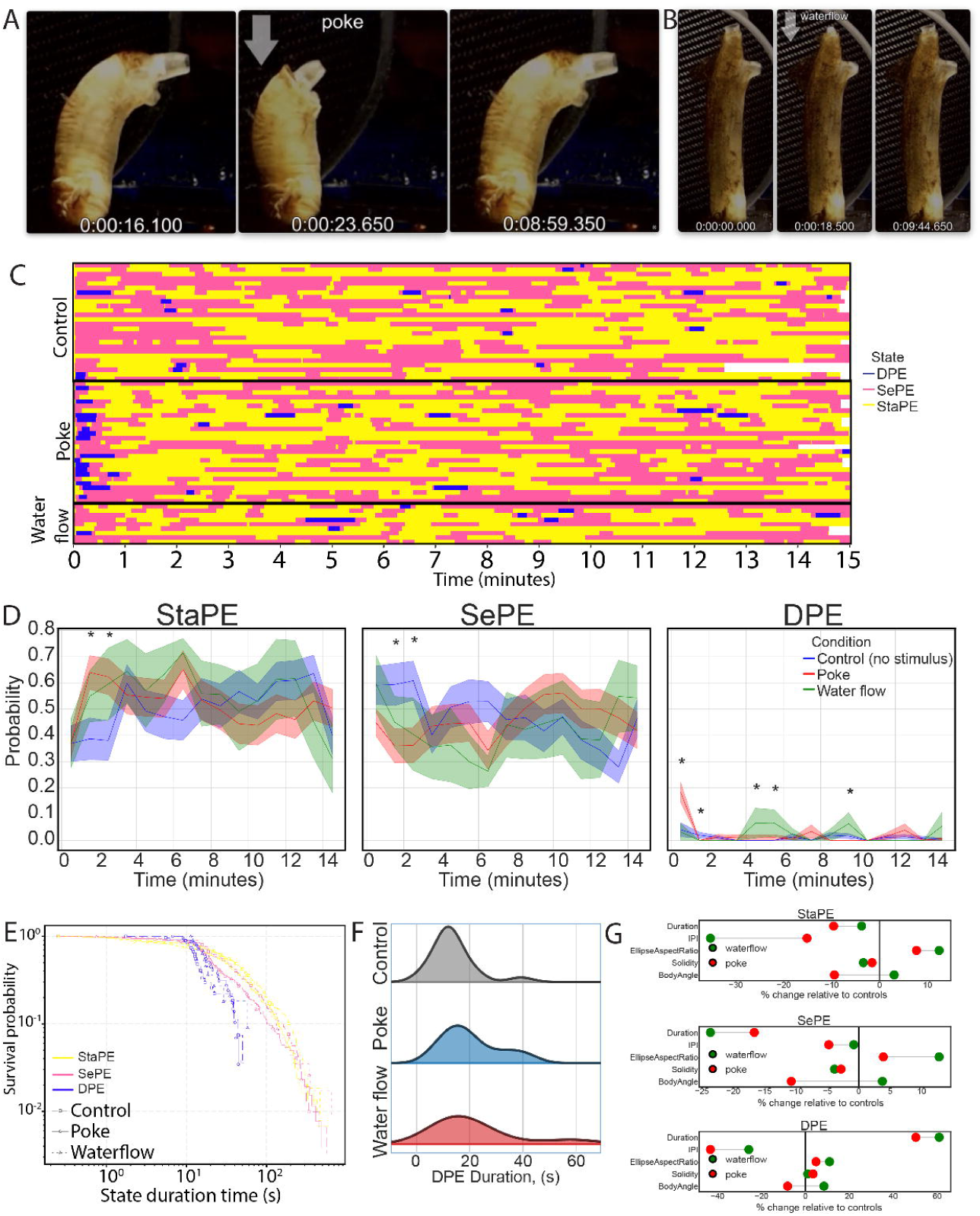
Mechanical stimuli elicit distinct behavioral responses. (A, B) Video frames from two *Ciona* adults responding to a (A) mechanical poke and (B) water flow (see Movies S2 and S3). In both cases the arrow indicates the time of stimulus application. (C) Ethograms of *Ciona* adults belonging to three groups (Controls i.e. no stimulus delivered, poke or water flow). Blue color corresponds to Dynamic postural engagement (DPE); pink corresponds to Selective postural engagement (SePE) and yellow to Static postural engagement (StaPE). (D) Line plots summarizing the probability of an adult Ciona being in one of the three states during the video recordings as a function of the stimulus used (no stimulus/control), poke and water flow. We show mean (thick lines) and (standard error of the mean) SEM shaded areas. For statistical analysis we used Kruskal-Wallis test (* p<0.05). (E) Quantification of the duration/persistence of each behavioral state as a function of stimulus used. (F) Distribution plots of the duration of DPE state events in animals that have not been exposed to a stimulus and animals exposed to mechanical poke and water flow. (G) Plots quantifying the percentage change relative to controls (no stimulus) of various postural and temporal features across the three different behavioral states.

In contrast to a poke stimulus, a water flow stimulus did not induce an immediate transition to a DPE state, suggesting that it is perceived by the animal’s nervous system as a qualitatively different stimulus to poke (Figure 2D). However, an unexpected observation was that that probability of animals entering a DPE state (mostly in the form of full contraction events) was significantly higher at 5 and 10 minutes after water flow stimulation, suggesting that this stimulus may elicit a periodic contraction of the animal whose biological significance is unclear (Figure 2D).

Inspecting the ethograms of unstimulated and stimulated animals we noticed that that the duration of individual StaPE, SePE and DPE events was different across the three treatments (Figure 2D). We found that in control unstimulated animals the static state (StaPE) was the most long-lasting (Figure 2E, yellow line with squares) with SePE state events having slightly shorter duration (Figure 2E, pink line with squares). The DPE state events were the most short-lived, which is reasonable given that this is an energetically costly state demanding the fast and recurrent contraction of multiple muscle sets along the animal’s body (Figure 2E, blue line with squares). We then asked whether mechanical stimulation in the form of poke or water flow altered the duration of individual events for each of the three states. The duration of individual events for postural quiescence and partial contraction was reduced in poke and water flow stimulated animals compared to the unstimulated control experiments (Figure 2E and 2G). However, DPE events had distributions that were shifted towards longer event lengths, showing a significant increase in event duration compared to unstimulated control events (Figure 2E-G). In addition, we quantified the interval between successive events of the same state (IPI). Static postural engagement (StaPE) and Dynamic postural engagement (DPE) states showed a significant decrease in the IPI in mechanically stimulated animals compared to unstimulated controls (Figure 2G). Two conditions were also different in transition state probabilities (Figure S2 E, F) and shape metrics (Figure S2 G-I). These surprising findings suggest that the nervous system of adult *Cionas* can modulate the duration and usage rate of these behavioral states in a sensory context-specific manner.

Amongst the postural features, the most striking effect we quantified was the ability of water flow stimulation to reduce the body angle of the animals across all behavioral states relative to control animals (Figure 2G). Interestingly, a poke stimulus had the opposite effect resulting in a slight increase in body angle (Figure 2G).

A distinguishing feature of SePE and DPE states in water flow stimulated animals is the oral siphon area relaxation speed which is much higher compared to non-stimulated and poke stimulated animals (Figure S2H and S2I). During StaPE periods oral siphon relaxation events are much faster for both water flow and poke (Figure S2G). Atrial siphon kinematics are significantly slower during the StaPE and SePE periods in water flow stimulated animals. Taken together our results suggest that the nervous system of *Ciona* adults can distinguish between the two qualitatively different sensory inputs and thus modulate accordingly the animal’s body posture.

### Five eigenciona shapes capture most of the postural variance of adult Ciona

While biologically interpretable features like amplitudes of trunk or siphon rim width opening amplitude, aspect ratio, body angle, dorsal/ventral ratio, and EFDs can describe postures with good accuracy, using these features comes at the price of very high dimensionality(*70*). We were thus interested in obtaining a simpler representation that described the range of postures that Ciona adults can attain as represented by the top ten harmonic (EFD) amplitudes without losing substantial information, since they explain over 95% of the total shape variance (Figure 3A; Figure S3A) (*71, 72*). We performed Principal Component Analysis (PCA) of these 10-dimensional vectors (amplitudes) and revealed that the first five components (here after described as eigencionas) explained over 87% of the variance (Figure 3A; Figure S3B). For any video frame, the body postures can be approximated as a linear combination of the five eigencionas: EC1 to EC5 (Figure 3A). Therefore, the coefficients of the five eigencionas were leveraged for further analysis as a simplified but accurate description of body postures (Figure 3B). Interestingly, while postural quiescence and partial contraction states showed relatively balanced contributions across all ECs, Dynamic postural engagement (DPE) were dominated by EC2 and EC3, corresponding to postures with contracted body and closed siphons (Figure 3A, 3C and 3D; Movie S4). To further capture the temporal dynamics of shape transitions, we also computed frame to frame derivatives of the 10 EFD amplitudes and performed PCA on these dynamic features (Figure S3E and S3F). The resulting dynamic principal components provided complementary information for state characterization, showing increasing dynamic contributions across most components from StaPE, SePE and DPE states (Figure 3E; Figure S3C-S3E).

**Figure 3.**
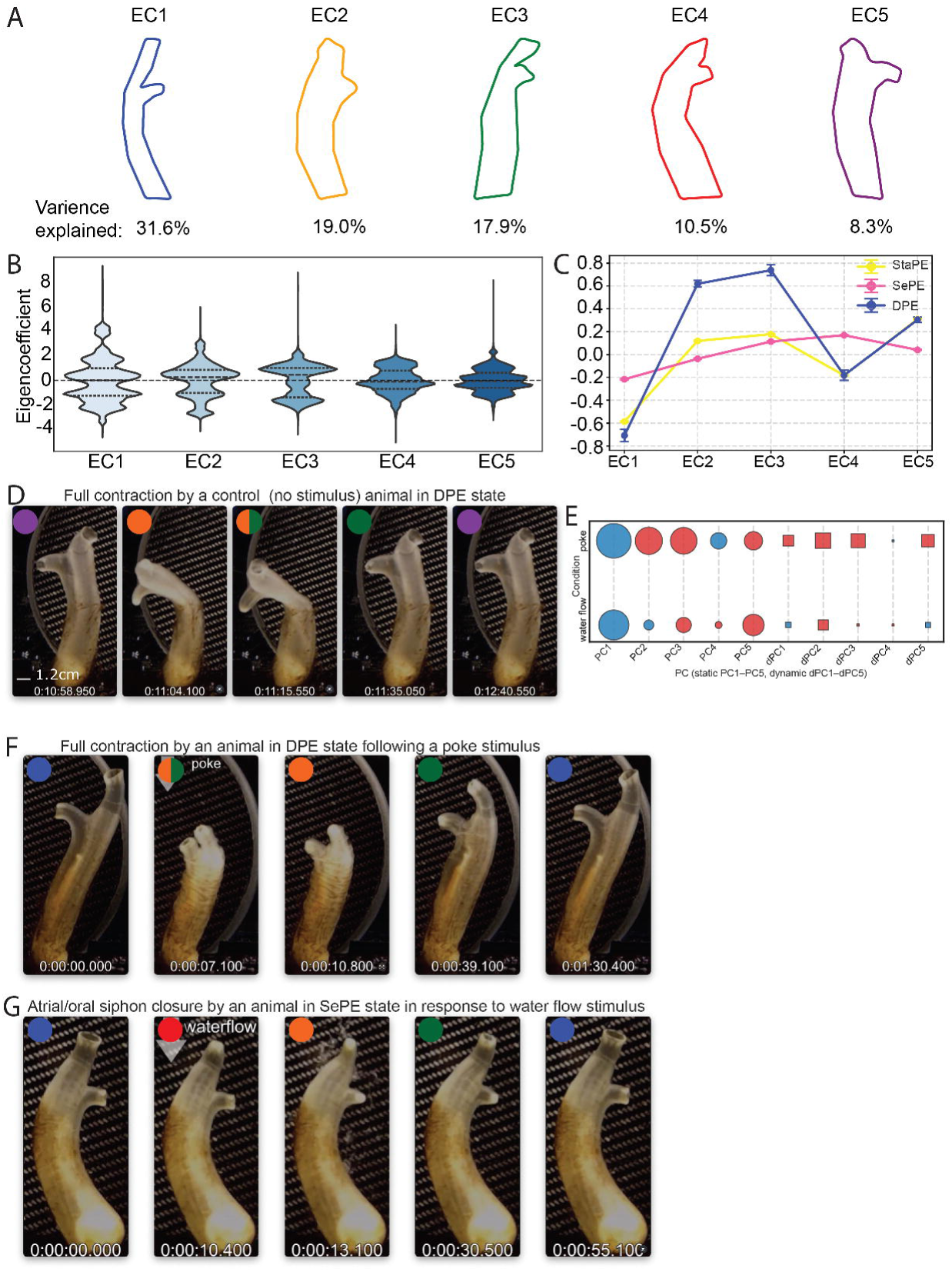
Five eigenciona shapes capture most of the postural variance of adult *Ciona*. (A) Visualization of the top 5 eigencionas obtained by an eigen decomposition of the covariance matrix, shown in descending order of the fraction of variance explained. (B) Distribution of the eigencoefficient values for all spontaneously behaving *Ciona* adults. (C) Eigenciona use summary by behavioral state. (D) Video frames from a *Ciona* adult exhibiting a spontaneous full contraction as they enter a DPE state. Coloring of the circles and semi-circles indicate the eigencionas that capture the postures of the *Ciona* in the specific video frames (see Movie S4). (E) Bubble grid chart showing the effects of poke and water flow stimuli in the use of static and dynamic eigencionas. Circles refer to static EigenCionas and boxes refer to dynamic eigencionas. Color indicates an increase (Red) or decrease (Blue) in eigenciona use, with the radius indicating the mean value. (F, G) Video frames of individual Cionas showing a response to a poke stimulus (F)and to a water flow stimulus (G) (see Movies S5 and S6). Color(s) of the circles and semi-circles indicate the eigencionas that capture the postures of the *Ciona* in the specific video frames.

To define shape responses to mechanical (poke) and hydrodynamic (water flow) stimuli in an unsupervised manner, we analyzed both static (PC1:PC5) and dynamic (dPC1:dPC5) components of body shape variation based on EFD amplitudes and compared them to spontaneous behavior (Figure 3E). In both conditions, PC1 captured the dominant static shape variation. Under poke, additional static components (PC2 and PC3) contributed substantially, reflecting stimulus-driven shape modulation, consistent with previous analyses highlighting the role of EC2 and EC3 when animals are found in a Dynamic postural engagement (DPE) state. In contrast, under water flow, static variation beyond PC1 was reduced. Similarly, dynamic shape changes under poke were prominent, particularly in dPC2 and dPC3, indicating active modulation of body shape in response to the mechanical stimulus (Figure 3E and 3F; Movies S5 and S6). In water flow, dynamic components were largely suppressed, suggesting more stable postures with limited shape modulation (Figure 3E and 3G). These findings show that poke stimuli evoke both static and dynamic shape changes, while water flow induces primarily static adjustments with limited dynamics.

### Low-dimensional spatiotemporal embedding reveals stereotyped actions in sessile adults

In earlier sections of our manuscript, we considered *Ciona* adult behavior as a sequence of discrete actions and states that we represented in ethograms. An assumption at the core of this approach is that *Ciona* adults perform behavioral actions in a way that allows for their discrete classification(*60*). However, we wanted to leverage a complementary approach aiming to reveal new structures in our behavioral data that may have been hard to detect initially by relying on human observations. To this end, we employed a spatiotemporal embedding approach(*19*) to reveal stereotyped behaviors exhibited by *Ciona* adults using unsupervised segmentation.

For this approach, we clustered shape trajectories based on PCA-transformed EFD amplitudes. We computed static, dynamic (derivative), and combined PCA spaces. Next, we obtained a lower dimensional “behavioral space” we employed a t-distributed stochastic embedding (tSNE), providing an embedding in which the local structure is retained. Subsequently this embedding which we can consider as the adult behavioral space, was sub-clustered using the DBSCAN algorithm (Figure 4A). We identified 18 clusters corresponding to different stereotyped behavioral modules (Figure 4B). Interestingly, this approach allowed us to identify stereotyped behavioral modules that are hard to discern by manual approaches. For example, cluster 10 corresponded to animals exhibiting a moderate withdrawal of the siphons towards the trunk, while their aperture remained open. On the other hand, cluster 6 corresponded to animals with elongated siphons whose aperture was closed. Another interesting example is cluster 1, where we observe a retraction of the closed atrial siphon but not the oral siphon. This is combined with a contraction of the circular trunk muscles leading to the reduction in body cross-section area.

**Figure 4.**
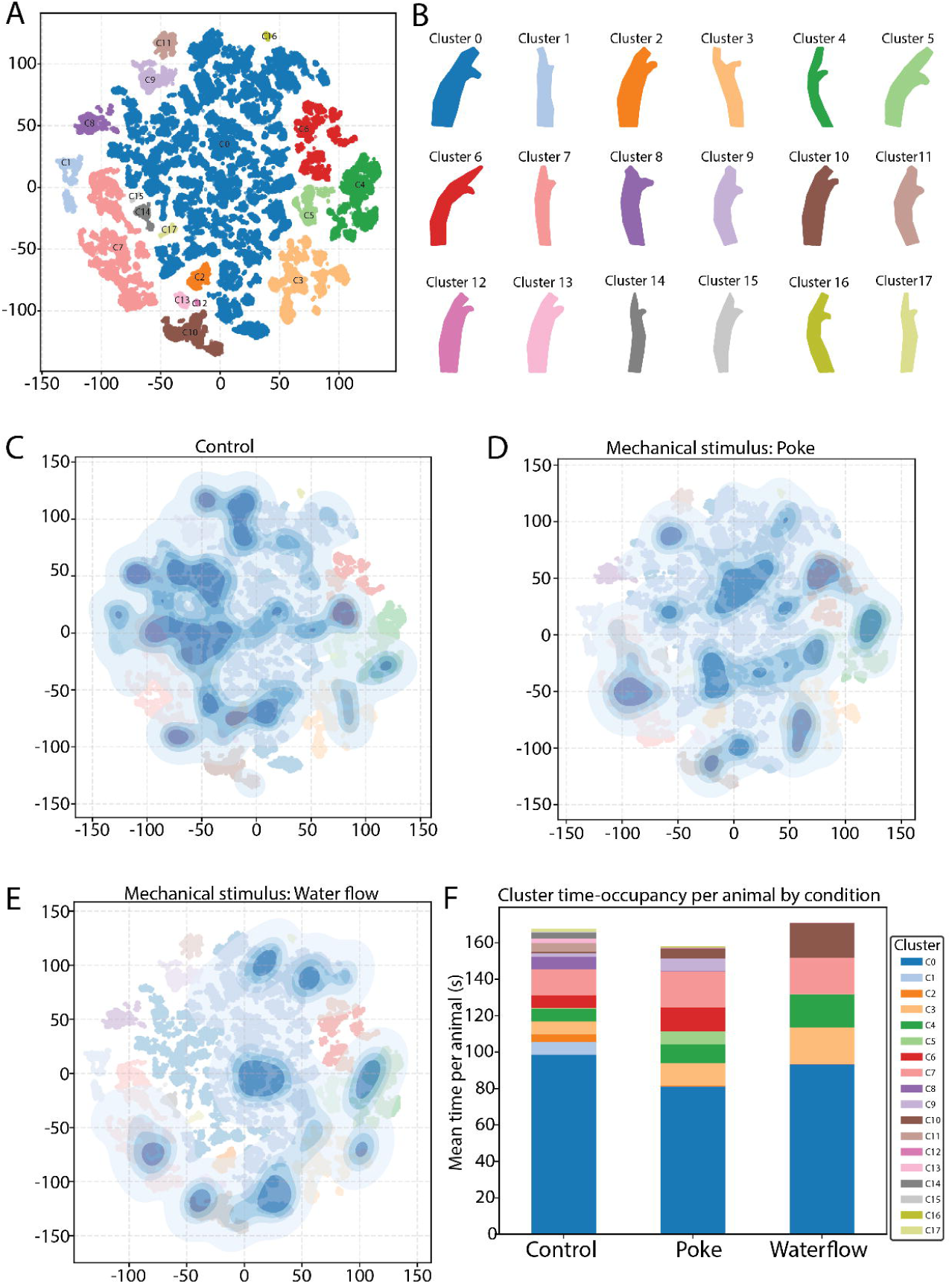
Low-dimensional spatiotemporal embedding reveals stereotyped actions in sessile adults. (A) t-SNE based 2D behavioral space of Ciona adults. The DBSCAN algorithm was used to identify clusters based on their density. (B) Examples of *Ciona* adult postures for each cluster. (C-E) KDE plots showing the change in the occupancy patterns of the 2D behavioral space across control (C), poke (D) and water flow (E) stimuli. The 18 DBSCAN clusters are also visible (opaque colors) under the KDE contours. (F) Stacked bar showing cluster time-occupancy per animal by condition.

We generated Kernel density estimation (KDE) plots for control unstimulated animals (spontaneous behaviors only) as well as for animals which were presented either with a poke or a water flow stimulus (Figure 4C-E). Far from being uniformly distributed across the behavioral space, the probability density contains several resolved local maxima. The locations of these maxima provide a potential representation for the stereotyped behaviors that the *Ciona* adults perform. Unstimulated control animals visited most regions of the behavioral space showing a large diversity of behavioral modes (Figure 4C). Animals stimulated with a mechanical poke showed fewer maxima indicating a more limited use of behaviors in response to mechanical poke perturbation (Figure 4D). Interestingly, local maxima were observed in clusters which are indicative of closing of both the atrial and oral siphons and/or a full body contraction (e.g. clusters 6 and 9), consistent with the strong response we see in the videos. Water flow probability density was concentrated in a few clusters relative to control and poke stimulated animals. These clusters represent behaviors where the atrial siphon is closed (e.g. cluster 4) or both siphons are open yet moderately withdrawn towards the trunk (e.g. cluster 10) consistent with the responses we have observed in the original movies.

Finally, we quantified the time spent by *Ciona* adults (per individual) on each cluster/ behavioral module. We found that animals across all conditions (control, poke and water flow stimulated) spent most of their time in cluster 0 followed by cluster 7, cluster 3 and cluster 4. These clusters corresponded primarily to relaxed postures with a fully relaxed/elongated trunk and either both or one of the siphons fully extended and with open aperture (Figure 4F). Beyond the four dominant behavioral clusters, the control animals exhibited a short amount of time in residual clusters of the behavioral space in contrast to mechanically stimulated animals which spend time longer periods of time in a smaller subset of clusters (Figure 4F). For example, animals exposed to a poke stimulus spent a longer time in behavioral clusters 6 and 9 compared to control and water flow stimulated animals, which are characteristic behaviors upon poking stimulus delivery (cluster 9) and following the poking stimulus (cluster 6). In the case of water flow stimulated animals dwell for longer in the behavior cluster 10 (Figure 4F). This is in line with what we have observed in the movies where a water flow stimulus elicits a withdrawal of the atrial and oral siphons towards the trunk but does not link to a contraction of the trunk.

## DISCUSSION

Here we present a quantitative framework for characterising the spontaneous, and stimuli evoked behaviors exhibited by adult *Ciona intestinalis* individuals. Compared to past behavioural studies on adult ascidians, this study benefits from cutting-edge tracking and analysis tools, leading to a data-rich and highly quantitative and detailed characterization of the behavioral repertoire of adult *Ciona intestinalis*. In our approach, posture information can be leveraged to detect subtle behaviors that would not have been discernible from mere qualitative observations or the use of other forms of tracking (e.g. centroid based). Thus, a major advancement of this work is the use of the animal contour (outline) which was extracted in a high-throughput and reliable manner by using a trained deep neural network(*63*). In this way we measured morphometric features such as siphon contraction and relaxation speeds, siphon aperture size and body angle. Importantly, we have derived a dimensionality reduced representation of adult *Ciona* postures (i.e. eigencionas). We demonstrated that five adult eigencionas can be combined in different fractions to reconstruct almost the entirety of behaving adult *Ciona* postures. This likely reflects fundamental constraints on *Ciona* adult behavior, where all behaviors of the adults explore different regions of the same shape space. This is a phenomenon observed in several organisms including *C. elegans* (*19, 73*), *D. melanogaster*(*74*) and *D. rerio*(*75, 76*). Having established a compact and common basis for representing *Ciona* shapes in two key phases of the biphasic lifecycle the larva (*19*) and adult stages (this study), future studies can leverage this resource in the context of high-throughput behavioral phenomics screens (e.g. to study the impact of anthropogenic pollutants such as neuroendocrine disruptors on marine organisms (*77*)) or detailed neurobiological studies. Past and ongoing efforts to map the developmental origin (*51, 78*) and innervation patterns of the adult nervous system (*59*) can now be combined with an in depth study of the adult postural dynamics.

In parallel to employing body part kinematics, postural dynamics and eigencionas, we have also implemented a spatiotemporal mapping approach similar to what has been employed in mapping complex behavioral responses in mice, flies and more recently in *Ciona* larvae(*79–81*). Here we present 18 annotated clusters of different behavioral modes that segregate to discrete regions of the behavioral space map. An important conclusion of our work is that the behavioral responses we observed in response to different mechanical cues (poke and water flow), do not result in global changes in the underlying spatiotemporal structure of *Ciona* adult behavioral space, but rather they arise from the selective use of behavioral modules and changes in the amount spent in each behavioral module. This suggests that the *Ciona* adult brain can select which individual behavioral modules it will use (and for how long it will use them) to generate responses to novel situations such as the presentation of mechanical stimuli.

In this study we made the surprising discovery that *Ciona* adults, which have long been described as passive filter feeders with a rather simple nervous system (*56, 59*), can exhibit different behavioral states. These states are distinguished from each other by various postural and motion features, including but not limited to the opening and closing speed and amplitude (aperture) of the oral and atrial siphons and the rate and extent of contraction of the trunk (i.e. how fast and how much they shrink and relax their body). Importantly, they can persist for different lengths of time and exhibit distinct transition probabilities. We have found that transitions between behavioral states can be induced by external stimuli (water flow and poke) but also they can occur probabilistically in the absence of a defined trigger stimulus (as seen in our control animals), suggesting that they can be internally generated(*82, 83*).

In other model organisms like *C. elegans* the duration of behavioral states is influenced by factors like sensory cues and the animal’s internal state(*68, 84, 85*). In the case of *Ciona* adults, we demonstrate that sensory cues can alter the duration of states and especially of the Dynamic Postural Engagement (DPE) state. Interestingly, a poke stimulus that induces a very strong and lasting DPE state is followed at a higher probability (compared to no stimulus condition) by StaPE state (i.e. a low activity state). While this may be a surprising observation in the first instance, behavioral challenges can cause animals to switch from active to passive strategies for coping with effort expenditure during stress(*86*). How might these states be modulated at the molecular and cellular levels? The oral and atrial siphons which show a strong modulation based on the behavioral state of the *Ciona* adult are innervated by a series of anterior and posterior peptidergic nerves (*59*). Given the richness of neuropeptides present in the adult nervous system(*87–90*), it is plausible that peptidergic volume transmission is involved in modulating the duration and features of the behavioral states exhibited by *Ciona* adults akin to what has been observed in other nervous systems across the tree of life(*91–95*).

How is it that adult *Ciona* can distinguish between these two different mechanical cues? One possibility is that *Ciona* adults use distinct populations of sensory cells to detect these different mechanical cues. *Ciona* is equipped with 75-100 cupular sense organs which are found in the lining of the atrial siphon (*42, 96*). The localization of the cupular organs close to the atrial siphon indicates that they may monitor water flow, however, to date we lack supportive functional evidence. Conversely, the oral siphon of *Ciona* adults is equipped with a row of secondary sensory cells which form the coronal organ(*97, 98*). In the colonial tunicate *B. schlosseri* these cells are thought to mediate the oral siphon stimulation test (an acute waterjet stimulus)(*99*), while in the solitary ascidian *Corella inflata* they respond to touch by a needle(*55*). Thus, it is conceivable, that the coronal organ is responsive to poke stimuli. The ability to distinguish water flow stimuli from other mechanical cues is of profound importance given that most sessile filter feeders depend on ambient water flow to replenish their food supplies and to remove waste products(*100*). Similarly the ability to detect acute mechanical stimuli (e.g. a poke) that may correspond to a benthic predator, such as sea urchin, crabs, sea stars and others is equally important for the survival of juvenile and adult ascidians(*101–104*).

To date, reports on the behavior of invertebrate sessile benthic organisms have been sparse and have mostly focused on *Poriferans* (i.e. sponges)(*30, 31*) and abyssal anemones(*105, 106*). One of these studies has used convolutional neural networks to study sponge behavior over time(*30*). Benthic communities are linked with various processes occurring at the sea surface, despite the vertical depth distance that can separate the two. Many studies have focused on highly motile fauna, while the sessile fauna of benthic environments has largely been ignored despite several observations that sessile organisms can show dynamic lifestyles showing both spontaneous and rhythmic behaviors(*30, 31*). We believe that our approach and investigation of behavioral states can be extended to other ascidian adults as well as other sessile benthic organisms like barnacles and articulated coralline algae.

Finally, from an evolutionary perspective our work on the behavior of *Ciona* adults allows us to get a glimpse of the possible behavioral repertoire of extant sessile epibenthic organisms. Our study may be particularly relevant for mid-Cambrian tunicates like *Megasiphon thylakos* which have both circular and longitudinal musculature like modern ascidians(*13*). It is thus conceivable that together with the establishment of the fundamental components of the modern tunicate body plan the behavioral traits of tunicates had already evolved almost 500Mya ago.

## RESOURCE AVAILABILITY

### Lead contact

Further information and requests for resources and reagents should be directed to and will be fulfilled by the lead contact, Marios Chatzigeorgiou

### Materials availability

This study did not generate any new materials.

### Data and code availability

- Raw video data have been deposited in Zenodo. The DOIs are: https://doi.org/10.5281/zenodo.16608643 https://doi.org/10.5281/zenodo.16614277
- The DeepLabCut based DNN trained to recognize landmarks on Ciona intestinalis adults has been deposited in Zenodo together with the training dataset. The DOI is: https://doi.org/10.5281/zenodo.16910611
- All original code will be deposited at Zenodo and will be publicly available as of the date of publication. Code to analyze behavioral data, is already available here: https://github.com/ChatzigeorgiouGroup/Ciona_adult_behavior
- Any additional information required to reanalyze the data reported in this paper will be made available from the lead contact upon request.

## ACKNOWLEDGMENTS

We would like to thank members of the Chatzigeorgiou lab for valuable feedback on the manuscript. We acknowledge funding from the Research Council of Norway: 339399 (M.C.) and 335582 (M.C.).

## AUTHOR CONTRIBUTIONS

Conceptualization: O.T., S.N., M.C.; Methodology: O.T., S.N., M.C.; Data curation: O.T., S.N.; Investigation: O.T., S.N.; Visualization: O.T., S.N.; Resources: M.C.; Funding acquisition: M.C.; Project administration: M.C.; Supervision: M.C.; Writing – original draft: O.T., S.N., M.C.; Writing – review and editing: O.T., S.N. M.C.

## EXPERIMENTAL MODEL AND STUDY PARTICIPANT DETAILS

We harvested wild adult *C. intestinalis* (*Ciona* intestinalis Type B) from the following sites: Døsjevika (60.342564, 5.114036), Kleppholmen (60.185215, 5.152785), Klokkarvik (60.222616, 5.157331), Liaskjeret (60.294301, 5.132846) and Toftøyna (60.482154, 4.950136), Sotra, Vestland, Norway. Wild *Cionas* were transferred to a purpose-built facility at the Michael Sars Centre, University of Bergen. We kept approximately 100 adults in 50L tanks with constant running filtered sea water (FSW) at 10 to 12°C and provided constant light and daily food supply. The diet included several species of diatoms and brown algae to increase egg production and reduce spontaneous spawning(*18*). Wild adults were allowed to acclimatize in the indoor facility for at least 24 h before behavioral recording.

## METHOD DETAILS

### Behavioral recording

On the day of recording, one or two adults were transferred to 54 L experimental glass tanks (60 cm × 30 cm) filled with FSW and 5 ml of algae mixture (1.4 ml *Chaetoceros calcitrans*, 1.4 ml *Rhinomonas*, 1.4 ml *Isochrysis sp*., 700 µl *Synechoccoccus sp*., from algal stock culture), and overhead lights (color temperature = 2700 K, Mira 26B dual flex arm, NANLITE, China) were provided. In the glass tank, the animal posterior region of the body (holdfast-end) was gently mounted vertically in a “mounting rack” located at the tank bottom. This mounting rack consisted of the top panel of a tube rack (Snap-Together canonical tube rack for 15 ml tubes, Heathrow Scientific, IL, USA) fixed on top of a plastic basket. The tube rack was covered with waterproof black spray paint to optimize the optical field. Additional weight was added to the basket allowing the rack to sink to the bottom of the tank as well as preventing movement during recording. The animals recorded were selected for their height and coverage of other organisms. The height of adult *Ciona* was 11.31 cm ± 2.62 cm after mounting. In cases where *Ciona* was covered by other organisms, such as algae or smaller *Ciona* that had settled on an adult, both oral and atrial siphon rims and cavities needed to be visible and differentiated from the lateral side. The choice of one or two animals recorded at once was based on their morphological curvature after mounting on the racks despite the provided overhead lights. Two animals were used if the angle of the mounted animals did not prevent full visualization of the neighboring animal during recording. The animals were mounted in the dorsoventral orientation according to the camera position for recording, allowing visualization of both siphons. The animals were recorded using Logitech HD Pro c920 (Logitech, Switzerland) controlled by Open Broadcaster Software (OBS) Studio (https://obsproject.com/). Data were recorded at 20 frames per second.

### Mechanical exposure assay

After transfer and mounting in the rack, adult *Ciona* was acclimated in the experimental tank for 15 min without any disturbances, followed by recording for 15-60 min in the absence of external stimuli to investigate individual variations in the behavioral repertoire (control). In the mechanical exposure assays, behavioral responses were recorded for 15 min in response to (i) poking the outer area of the oral siphon with a glass Pasteur pipette and (ii) a water current by fluid flow (28 ml/s) initiated by introducing FSW into the tank. After recording, the adult was removed from the tank, stroked twice with a glass Pasteur pipette from posterior to anterior end, and gently wiped with paper to remove remnant surface water, before their wet weight was logged.

### DeepLabCut model

The adult *Ciona* DLC model was trained to track 18 body points from the dorsal to the ventral and anterior to posterior positions based on external morphological characteristics, including the rims and cavities of the oral and atrial siphons, and along the trunk region. The adult DLC model was trained using 20 frames from 22 randomly selected videos (440 frames in total) for 200.000 iterations and three shuffles (other parameters were kept at default) with the ResNet50 ImageNet pre-trained weight architecture as the selected network architecture. 10 outlier frames from 34 videos were corrected following 100.000 iterations and three shuffles. The DLC model was characterized by train and test errors of 3.15 and 3.91 pixels, and 3.14 and 3.91 with 0.6 p-cutoff value. Mm-to-pixel conversion ratio was determined *in silico* to 0.254 mm/pixel. Data was generated for all videos, and the filtered tracking data were exported as CSV files and used for extracting features. Filtered coordinate tracking data (DeepLabCut outputs) were loaded from individual .csv files and merged after removal of confidence (likelihood) columns. Each animal’s body part coordinates (x and y) were split, resampled (1:5 temporal downsampling resulted in 4 Hz signal), and annotated based on experimental condition (control, water flow, or poke).

### Extraction of supervised behavioral metrics

We computed six quantitative time series per animal: whole-body area (via 2D polygon surface), oral and atrial siphon opening widths (Euclidean distance between respective dorsal and ventral points), corresponding areas (defined by four-point contours), and a mantle area (polygon excluding siphon points). Only traces with ≥20% non-NaN data for all metrics were retained. Metric values were normalized by median, outliers filtered, missing data segments (≤20 frames) were interpolated. We applied seasonal decomposition (additive model, 50-frame periodicity) to isolate contraction trends from each metric. Trends were z-scored, extrapolated at the start (25-frame linear fit), and interpolated (≤200-frame gap fill). Initial and terminal valid points were marked to prevent boundary artifacts.

Contraction event detection on z-scored trends was performed using a smoothed signal (Savitzky-Golay filter), identifying contraction events via prominence based peak detection (minimum prominence: 0.7; minimum width: 5 frames). Detected events were categorized as ‘DPE’ or ‘SePE’ events based on relative peak amplitude with 70% of maximum peak threshold. StaPE periods were defined as time intervals with sub-threshold activity across three most informative metrics (mantle, oral siphon opening width, and atrial siphon opening width).

To quantify behavioral dynamics in Ciona, we first identified three recurrent states: DPE, SePE, and StaPE, by aligning frame by frame annotations for three informative metrics. Short transient states (<2 seconds) were merged with adjacent states to reduce noise. From this cleaned sequence, state events were extracted per animal and condition, with start/end frames and durations calculated over a 15-minute recording window. We computed inter event intervals, event durations and event frequencies (events/min) per state, and visualized state transitions using normalized chord diagrams. For each contraction event, we also quantified amplitude and the speeds of contraction and relaxation across morphometric metrics, after converting pixel measurements to micrometers.

To quantify how body shape changes across behavioral states, we extracted postures (shape outline) for each animal during DPE, SePE, and StaPE, states. From these outlines we calculated three shape metrics: ellipse aspect ratio which reflects body elongation and was estimated by fitting an ellipse to the body shape and taking the ratio of its long to short axes; solidity, which measures how tightly the body outline fits its convex hull and captures body curvature or folding; and body angle, defined as the orientation of the main body axis relative to the horizontal, indicating dorsal tilt. To account for differences in individual body size or orientation, each metric was normalized to the each animal’s median across all three states. All metrics were averaged per individual and condition before plotting.

### Extraction of unsupervised behavior features

We used the outline coordinates of each animal at each time point and converted them into closed polygons. Invalid shapes (e.g., incomplete or geometrically impossible contours) were excluded. We used Elliptic Fourier Descriptors (EFDs) to represent each outline as a set of harmonics that capture shape features at increasing detail. Fourier analysis was conducted using the pyefd package (https://github.com/hbldh/pyefd), following the Kuhl & Giardina (1982)(*70*). Specifically, the elliptic_fourier_descriptors function was employed with a default order of 30, generating 120 coefficients (30 orders × 4 coefficients per order). This choice was confirmed visually by comparing original outlines with EFD reconstructions (error <1%). To ensure consistency, each shape was normalized for size and aligned to correct for left right mirroring. A data-driven scan across samples showed that 10 harmonics were sufficient to capture ≥99.9% of the overall shape information, and this value was used throughout for quantification.

We applied principal component analysis (PCA) to a 10-component amplitude spectrum to identify major modes of variation for static shape analysis. To include dynamics, we also computed how these amplitudes changed over time performing separate PCA.

These 10-dimensional amplitude vectors were then z-scored and submitted to Principal Component Analysis (PCA) using scikit-learn (Pedregosa et al., 2011)(*107*). We retained the smallest number of PCs that explained ≥ 85 % of the variance. The resulting PC1–PC5 scores therefore serve as our unsupervised “eigen shape” metrics or “eigen cionas”, capturing 85–90 % of the observed outline variability. To verify interpretability, eigen shapes (principal components) were visualized and confirmed by reconstructing real contours associated with extreme PC scores for each animal and behavior state. Downstream analyses (state-classification, statistical comparisons) were performed on these five dimensions /“eigen cionas”.

For the shape space embedding and to visualize relationships between frames based on body shape, we used the first five principal components of shape (static PCs) derived from three feature spaces. Each PC matrix was embedded into two dimensions using t-distributed stochastic neighbor embedding (t-SNE; openTSNE, perplexity = 150). Clusters in t-SNE space were identified using DBSCAN (ε = 6, min_samples = 50). Results were visualized by behavioral state and condition.

We also applied a wavelet transform to the EFD time series to extract multi-scale temporal features, summarizing how different shape components fluctuated across time and behavior states identified by supervised approach.

We used Hidden-Markov modelling for long-range shape dynamics. EFD time series were divided into 20-s windows with 50 % overlap (frame interval = 0.25 s; 80 frames per window, 40-frame step). Within each window, we computed a Morlet continuous wavelet transform (scales 1–50) for every EFD coefficient and averaged the absolute coefficients across time, yielding a 1500-element vector (30 harmonics × 50 scales). Features were z-scored and reduced to five principal components before training a five-state Gaussian hidden-Markov model (full covariance, 300 EM iterations). The model was fitted on control animals only and then used to decode state sequences for every window.

### Statistical comparisons

To make comparisons between states and conditions were performed using pairwise Mann Whitney U tests.

## REFERENCES

1. M. L. Droser, J. G. Gehling, The advent of animals: The view from the Ediacaran. Proc Natl Acad Sci U S A 112, 4865–4870 (2015).

2. G. M. Narbonne, THE EDIACARA BIOTA: Neoproterozoic Origin of Animals and Their Ecosystems. Annual Review of Earth and Planetary Sciences 33, 421–442 (2005).

3. Z. Yin et al., Sponge grade body fossil with cellular resolution dating 60 Myr before the Cambrian. Proc Natl Acad Sci U S A 112, E1453–1460 (2015).

4. M. L. Droser, J. G. Gehling, Synchronous Aggregate Growth in an Abundant New Ediacaran Tubular Organism. Science 319, 1660–1662 (2008).

5. I. A. Rahman, S. A. F. Darroch, R. A. Racicot, M. Laflamme, Suspension feeding in the enigmatic Ediacaran organism Tribrachidium demonstrates complexity of Neoproterozoic ecosystems. Sci Adv 1, e1500800.

6. J. R. Paterson, J. G. Gehling, M. L. Droser, R. D. Bicknell, Rheotaxis in the Ediacaran epibenthic organism Parvancorina from South Australia. Sci Rep 7, 45539 (2017).

7. J. Peng, L. E. Babcock, Y. Zhao, P. Wang, R. Yang, Cambrian Sphenothallus from Guizhou Province, China: early sessile predators. Palaeogeography, Palaeoclimatology, Palaeoecology 220, 119–127 (2005).

8. T. J. Kloss, S. Q. Dornbos, J. Chen, Substrate adaptations of sessile benthic metazoans during the Cambrian radiation. Paleobiology 41, 342–352 (2015).

9. Y. Zhao et al., Cambrian Sessile, Suspension Feeding Stem-Group Ctenophores and Evolution of the Comb Jelly Body Plan. Curr Biol 29, 1112–1125 e1112 (2019).

10. K. A. Kolesnikov, J. P. Botting, A. Y. Ivantsov, A. Y. Zhuravlev, New early Cambrian sponges of the Siberian platform and the origins of spiculate crown-group demosponges. Papers in Palaeontology 10, (2024).

11. B. Pan et al., Early Cambrian organophosphatic brachiopods from the Xinji Formation, at Shuiyu section, Shanxi Province, North China. Palaeoworld 29, 512–533 (2020).

12. H. Yun et al., Adaptive specialization of a unique sponge body from the Cambrian Qingjiang biota. Proc Biol Sci 289, 20220804 (2022).

13. K. Nanglu, R. Lerosey-Aubril, J. C. Weaver, J. Ortega-Hernandez, A mid-Cambrian tunicate and the deep origin of the ascidiacean body plan. Nat Commun 14, 3832 (2023).

14. J.-Y. Chen et al., The first tunicate from the Early Cambrian of South China. Proceedings of the National Academy of Sciences 100, 8314–8318 (2003).

15. V. Laudet, The Origins and Evolution of Vertebrate Metamorphosis. Current Biology 21, R726–R737 (2011).

16. Y.-B. Shi, in Current Topics in Developmental Biology, Y.-B. Shi, Ed. (Academic Press, 2013), vol. 103, pp. xv-xvi.

17. O. Tolstenkov et al., The cellular and behavioral blueprints of chordate rheotaxis. bioRxiv, 2025.2003.2022.644710 (2025).

18. J. Rudolf, D. Dondorp, L. Canon, S. Tieo, M. Chatzigeorgiou, Automated behavioural analysis reveals the basic behavioural repertoire of the urochordate Ciona intestinalis. Sci Rep 9, 2416 (2019).

19. A. Athira, D. Dondorp, J. Rudolf, O. Peytral, M. Chatzigeorgiou, Comprehensive analysis of locomotion dynamics in the protochordate Ciona intestinalis reveals how neuromodulators flexibly shape its behavioral repertoire. PLoS Biol 20, e3001744 (2022).

20. R. R. Strathmann, Larval feeding in echinoderms. Am Zool 15, 717–730 (1975).

21. R. R. Strathmann, T. L. Jahn, J. R. Fonseca, Suspension feeding by marine invertebrate larvae: clearance of particles by ciliated bands of a rotifer, pluteus, and trochophore. The Biological Bulletin 142, 505–519 (1972).

22. R. R. Strathmann, Time and Extent of Ciliary Response to Particles in a Non-Filtering Feeding Mechanism. The Biological Bulletin 212, 93–103 (2007).

23. S. Whalan, The role of photobehaviour in sponge larval dispersal and settlement. PLoS One 18, e0287989 (2023).

24. S. P. Leys et al., Sponge Behavior and the Chemical Basis of Responses: A Post-Genomic View. Integr Comp Biol 59, 751–764 (2019).

25. L. A. Bezares Calderón, R. Shahidi, G. Jékely, Mechanism of barotaxis in marine zooplankton. eLife 13, RP94306 (2024).

26. L. A. Bezares-Calderón et al., Neural circuitry of a polycystin-mediated hydrodynamic startle response for predator avoidance. eLife 7, e36262 (2018).

27. M. A. Tosches, D. Bucher, P. Vopalensky, D. Arendt, Melatonin signaling controls circadian swimming behavior in marine zooplankton. Cell 159, 46–57 (2014).

28. G. Jekely et al., Mechanism of phototaxis in marine zooplankton. Nature 456, 395–399 (2008).

29. J. Hoyer et al., Polymodal sensory perception drives settlement and metamorphosis of Ciona larvae. Curr Biol 34, 1168–1182 e1167 (2024).

30. D. Harrison et al., Machine Learning Applications of Convolutional Neural Networks and Unet Architecture to Predict and Classify Demosponge Behavior. Water 13, (2021).

31. A. S. Kahn, C. W. Pennelly, P. R. McGill, S. P. Leys, Behaviors of sessile benthic animals in the abyssal northeast Pacific Ocean. Deep Sea Research Part II: Topical Studies in Oceanography 173, (2020).

32. F. Delsuc, H. Brinkmann, D. Chourrout, H. Philippe, Tunicates and not cephalochordates are the closest living relatives of vertebrates. Nature 439, 965–968 (2006).

33. Z. Liang, J. Hoyer, M. Chatzigeorgiou, Anoctamins mediate polymodal sensory perception and larval metamorphosis in a non-vertebrate chordate. Cell Reports 44, (2025).

34. H. Wada, H. Saiga, N. Satoh, P. W. Holland, Tripartite organization of the ancestral chordate brain and the antiquity of placodes: insights from ascidian Pax-2/5/8, Hox and Otx genes. Development 125, 1113–1122 (1998).

35. L. Z. Holland, N. D. Holland, Chordate origins of the vertebrate central nervous system. Current Opinion in Neurobiology 9, 596–602 (1999).

36. I. A. Meinertzhagen, P. Lemaire, Y. Okamura, The neurobiology of the ascidian tadpole larva: recent developments in an ancient chordate. Annu Rev Neurosci 27, 453–485 (2004).

37. K. Ryan, Z. Lu, I. A. Meinertzhagen, The CNS connectome of a tadpole larva of Ciona intestinalis (L.) highlights sidedness in the brain of a chordate sibling. Elife 5, (2016).

38. A. Stolfi, M. Levine, Neuronal subtype specification in the spinal cord of a protovertebrate. Development 138, 995–1004 (2011).

39. A. Stolfi, E. Wagner, J. M. Taliaferro, S. Chou, M. Levine, Neural tube patterning by Ephrin, FGF and Notch signaling relays. Development 138, 5429–5439 (2011).

40. K. S. Imai, A. Stolfi, M. Levine, Y. Satou, Gene regulatory networks underlying the compartmentalization of the Ciona central nervous system. Development 136, 285–293 (2009).

41. S. Gibboney et al., Effector gene expression underlying neuron subtype-specific traits in the Motor Ganglion of Ciona. Dev Biol 458, 52–63 (2020).

42. C. Anselmi, G. K. Fuller, A. Stolfi, A. K. Groves, L. Manni, Sensory cells in tunicates: insights into mechanoreceptor evolution. Front Cell Dev Biol 12, 1359207 (2024).

43. K. Kim, K. M. Piekarz, A. Stolfi, A gene regulatory network for specification and morphogenesis of a Mauthner Cell homolog in non-vertebrate chordates. Dev Biol 522, 51–63 (2025).

44. K. M. Piekarz, A. Stolfi, Development and circuitry of the tunicate larval Motor Ganglion, a putative hindbrain/spinal cord homolog. J Exp Zool B Mol Dev Evol 342, 200–211 (2024).

45. S. Chiba, A. Sasaki, A. Nakayama, K. Takamura, N. Satoh, Development of Ciona intestinalis juveniles (through 2nd ascidian stage). Zoolog Sci 21, 285–298 (2004).

46. A. Nakayama-Ishimura, J. P. Chambon, T. Horie, N. Satoh, Y. Sasakura, Delineating metamorphic pathways in the ascidian Ciona intestinalis. Dev Biol 326, 357–367 (2009).

47. Y. Sasakura, A. Hozumi, Formation of adult organs through metamorphosis in ascidians. Wiley Interdiscip Rev Dev Biol 7, (2018).

48. H. D. Dufour et al., Precraniate origin of cranial motoneurons. Proc Natl Acad Sci U S A 103, 8727–8732 (2006).

49. T. Horie et al., Ependymal cells of chordate larvae are stem-like cells that form the adult nervous system. Nature 469, 525–528 (2011).

50. F. Razy-Krajka et al., Collier/OLF/EBF-dependent transcriptional dynamics control pharyngeal muscle specification from primed cardiopharyngeal progenitors. Dev Cell 29, 263–276 (2014).

51. E. D. Gigante et al., Specification and survival of post-metamorphic branchiomeric neurons in a non-vertebrate chordate. Development 151, (2024).

52. S. Hecht, The physiology of Ascidia atra Lesueur. I. General physiology. J Exp Zool 25, 229–259 (1918).

53. S. Hecht, The physiology of Ascidia atra Lesueur. II. Sensory physiology. J Exp Zool 25, 261–299 (1918).

54. E. C. Day, The physiology of the nervous system of the tunicate. I. The relation of the nerve ganglion to sensory responses. J Exp Zool 28, 307–335 (1919).

55. G. O. Mackie, P. Burighel, F. Caicci, L. Manni, Innervation of ascidian siphons and their responses to stimulation. Canadian Journal of Zoology 84, 1146–1162 (2006).

56. G. O. Mackie, P. Burighel, The nervous system in adult tunicates: current research directions. Canadian Journal of Zoology 83, 151–183 (2005).

57. K. Braun, T. Stach, Morphology and evolution of the central nervous system in adult tunicates. Journal of Zoological Systematics and Evolutionary Research 57, 323–344 (2018).

58. C. Dahlberg et al., Refining the Ciona intestinalis model of central nervous system regeneration. PLoS One 4, e4458 (2009).

59. T. Osugi, Y. Sasakura, H. Satake, The nervous system of the adult ascidian Ciona intestinalis Type A (Ciona robusta): Insights from transgenic animal models. PLoS One 12, e0180227 (2017).

60. A. E. X. Brown, B. de Bivort, Ethology as a physical science. Nature Physics 14, 653–657 (2018).

61. S. E. Egnor, K. Branson, Computational Analysis of Behavior. Annu Rev Neurosci 39, 217–236 (2016).

62. T. D. Pereira, J. W. Shaevitz, M. Murthy, Quantifying behavior to understand the brain. Nature Neuroscience 23, 1537–1549 (2020).

63. J. Lauer et al., Multi-animal pose estimation, identification and tracking with DeepLabCut. Nat Methods 19, 496–504 (2022).

64. A. Mathis et al., DeepLabCut: markerless pose estimation of user-defined body parts with deep learning. Nat Neurosci 21, 1281–1289 (2018).

65. T. D. Pereira et al., SLEAP: A deep learning system for multi-animal pose tracking. Nat Methods 19, 486–495 (2022).

66. J. M. Graving et al., DeepPoseKit, a software toolkit for fast and robust animal pose estimation using deep learning. Elife 8, (2019).

67. J. J. Sun et al., Self-Supervised Keypoint Discovery in Behavioral Videos. Proc IEEE Comput Soc Conf Comput Vis Pattern Recognit 2022, 2161–2170 (2022).

68. S. W. Flavell, D. M. Raizen, Y. J. You, Behavioral States. Genetics 216, 315–332 (2020).

69. S. W. Flavell, N. Gogolla, M. Lovett-Barron, M. Zelikowsky, The emergence and influence of internal states. Neuron 110, 2545–2570 (2022).

70. F. P. Kuhl, C. R. Giardina, Elliptic Fourier features of a closed contour. Computer Graphics and Image Processing 18, 236–258 (1982).

71. G. J. Stephens, B. Johnson-Kerner, W. Bialek, W. S. Ryu, Dimensionality and dynamics in the behavior of C. elegans. PLoS Comput Biol 4, e1000028 (2008).

72. G. J. Stephens, L. C. Osborne, W. Bialek, Searching for simplicity in the analysis of neurons and behavior. Proc Natl Acad Sci U S A 108 **Suppl 3**, 15565–15571 (2011).

73. A. E. Brown, E. I. Yemini, L. J. Grundy, T. Jucikas, W. R. Schafer, A dictionary of behavioral motifs reveals clusters of genes affecting Caenorhabditis elegans locomotion. Proc Natl Acad Sci U S A 110, 791–796 (2013).

74. B. Szigeti, A. Deogade, B. Webb, Searching for motifs in the behaviour of larval Drosophila melanogaster and Caenorhabditis elegans reveals continuity between behavioural states. J R Soc Interface 12, 20150899 (2015).

75. K. Girdhar, M. Gruebele, Y. R. Chemla, The Behavioral Space of Zebrafish Locomotion and Its Neural Network Analog. PLoS One 10, e0128668 (2015).

76. D. S. Mearns, J. C. Donovan, A. M. Fernandes, J. L. Semmelhack, H. Baier, Deconstructing Hunting Behavior Reveals a Tightly Coupled Stimulus-Response Loop. Curr Biol 30, 54–69 e59 (2020).

77. J. Beyer, Y. Song, A. Lillicrap, S. Rodriguez-Satizabal, M. Chatzigeorgiou, Ciona spp. and ascidians as bioindicator organisms for evaluating effects of endocrine disrupting chemicals: A discussion paper. Mar Environ Res 191, 106170 (2023).

78. L. G. Todorov, K. Oonuma, T. G. Kusakabe, M. S. Levine, L. A. Lemaire, Neural crest lineage in the protovertebrate model Ciona. Nature 635, 912–916 (2024).

79. G. J. Berman, D. M. Choi, W. Bialek, J. W. Shaevitz, Mapping the stereotyped behaviour of freely moving fruit flies. Journal of The Royal Society Interface 11, 20140672–20140672 (2014).

80. J. Cande et al., Optogenetic dissection of descending behavioral control in Drosophila. Elife 7, (2018).

81. J. D. Marshall et al., Continuous Whole-Body 3D Kinematic Recordings across the Rodent Behavioral Repertoire. Neuron 109, 420–437 e428 (2021).

82. E. H. Goulding et al., A robust automated system elucidates mouse home cage behavioral structure. P Natl Acad Sci USA 105, 20575–20582 (2008).

83. J. R. Martin, R. Ernst, M. Heisenberg, Temporal pattern of locomotor activity in Drosophila melanogaster. Journal of Comparative Physiology A 184, 73–84 (1999).

84. N. Ji et al., A neural circuit for flexible control of persistent behavioral states. eLife 10, e62889 (2021).

85. S. W. Flavell, M. Oren-Suissa, S. Stern, Sources of behavioral variability in C. elegans: Sex differences, individuality, and internal states. Curr Opin Neurobiol 91, 102984 (2025).

86. A. S. Andalman et al., Neuronal Dynamics Regulating Brain and Behavioral State Transitions. Cell 177, 970–985 e920 (2019).

87. H. Satake et al., Tachykinin and tachykinin receptor of an ascidian, Ciona intestinalis: evolutionary origin of the vertebrate tachykinin family. J Biol Chem 279, 53798–53805 (2004).

88. T. Kawada, T. Sekiguchi, T. Sakai, M. Aoyama, H. Satake, Neuropeptides, hormone peptides, and their receptors in Ciona intestinalis: an update. Zoolog Sci 27, 134–153 (2010).

89. T. Kawada et al., Peptidomic analysis of the central nervous system of the protochordate, Ciona intestinalis: homologs and prototypes of vertebrate peptides and novel peptides. Endocrinology 152, 2416–2427 (2011).

90. S. Matsubara et al., The significance of Ciona intestinalis as a stem organism in integrative studies of functional evolution of the chordate endocrine, neuroendocrine, and nervous systems. Gen Comp Endocrinol 227, 101–108 (2016).

91. G. Jekely et al., The long and the short of it - a perspective on peptidergic regulation of circuits and behaviour. J Exp Biol 221, (2018).

92. M. P. Nusbaum, D. M. Blitz, E. Marder, Functional consequences of neuropeptide and small-molecule co-transmission. Nat Rev Neurosci 18, 389–403 (2017).

93. E. Svensson et al., General Principles of Neuronal Co-transmission: Insights From Multiple Model Systems. Front Neural Circuits 12, 117 (2018).

94. P. H. Taghert, M. N. Nitabach, Peptide neuromodulation in invertebrate model systems. Neuron 76, 82–97 (2012).

95. U. S. Bhat, N. Shahi, S. Surendran, K. Babu, Neuropeptides and Behaviors: How Small Peptides Regulate Nervous System Function and Behavioral Outputs. Front Mol Neurosci 14, 786471 (2021).

96. Q. Bone, K. P. Ryan, Cupular Sense-Organs in Ciona (Tunicata-Ascidiacea). J Zool 186, 417–429 (1978).

97. F. Rigon, F. Gasparini, S. M. Shimeld, S. Candiani, L. Manni, Developmental signature, synaptic connectivity and neurotransmission are conserved between vertebrate hair cells and tunicate coronal cells. J Comp Neurol 526, 957–971 (2018).

98. P. Burighel, F. Caicci, L. Manni, Hair cells in non-vertebrate models: lower chordates and molluscs. Hear Res 273, 14–24 (2011).

99. C. Anselmi et al., Two distinct evolutionary conserved neural degeneration pathways characterized in a colonial chordate. Proc Natl Acad Sci U S A 119, e2203032119 (2022).

100. C. M. Young, L. F. Braithwaite, Orientation and Current-Induced Flow in the Stalked Ascidian Styela-Montereyensis. Biol Bull 159, 428–440 (1980).

101. C. B. Giachetti, N. Battini, K. L. Castro, E. Schwindt, The smaller, the most delicious: Differences on vulnerability to predation between juvenile and adult of invasive ascidians. Estuarine, Coastal and Shelf Science 268, (2022).

102. C. P. Dumont, C. F. Gaymer, M. Thiel, Predation contributes to invasion resistance of benthic communities against the non-indigenous tunicate Ciona intestinalis. Biological Invasions 13, 2023–2034 (2011).

103. A. Epelbaum, C. M. Pearce, D. J. Barker, A. Paulson, T. W. Therriault, Susceptibility of non-indigenous ascidian species in British Columbia (Canada) to invertebrate predation. Marine Biology 156, 1311–1320 (2009).

104. R. W. Osman, R. B. Whitlatch, The control of the development of a marine benthic community by predation on recruits. Journal of Experimental Marine Biology and Ecology 311, 117–145 (2004).

105. J. M. Durden, B. J. Bett, H. A. Ruhl, The hemisessile lifestyle and feeding strategies of Iosactis vagabunda (Actiniaria, Iosactiidae), a dominant megafaunal species of the Porcupine Abyssal Plain. Deep Sea Research Part I: Oceanographic Research Papers 102, 72–77 (2015).

106. R. Lampitt, G. Paterson, The feeding-behavior of an abyssal sea-anemone from insitu time-lapse photographs and trawl samples. Oceanologica Acta 10, 455–461 (1987).

107. F. Pedregosa et al., Scikit-learn: Machine Learning in Python. J. Mach. Learn. Res. 12, 2825–2830 (2011).

