## Supplementary figures and images for "The evolution of embodied postural dynamics underlies behavioral diversity in benthic sessile chordate"

### Supplemental Figure 1

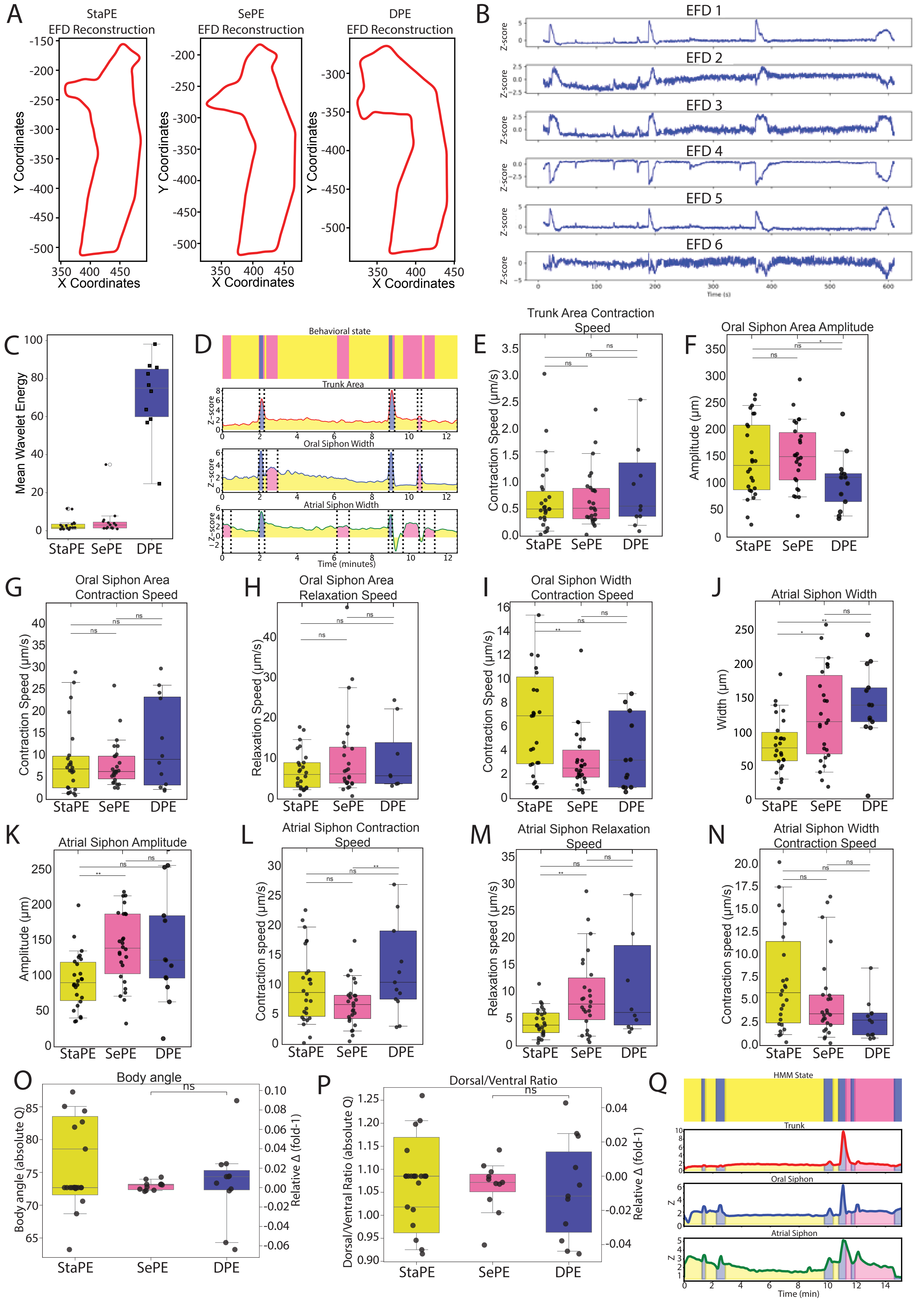

### Supplemental Figure 2

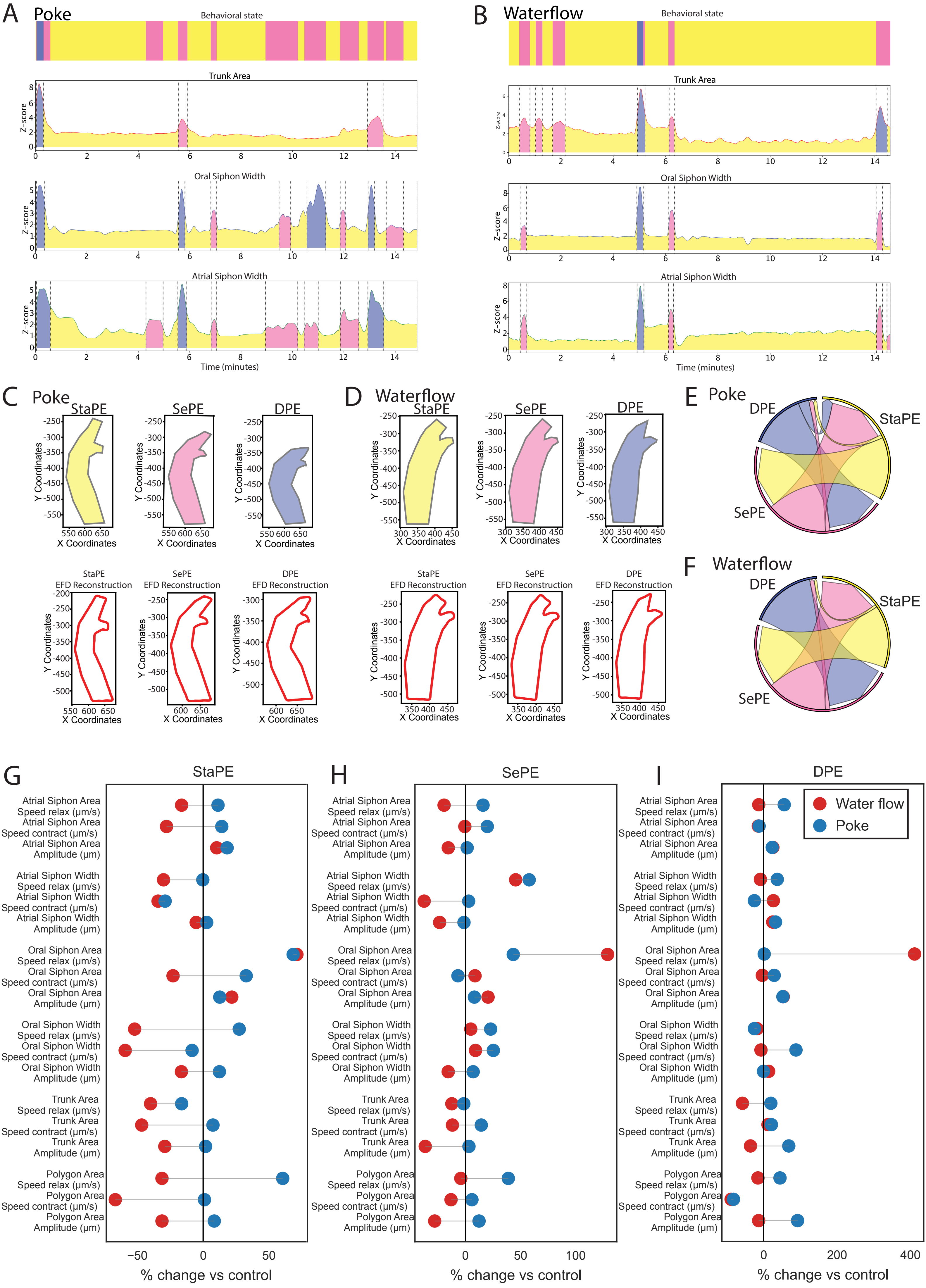

### Supplemental Figure 3

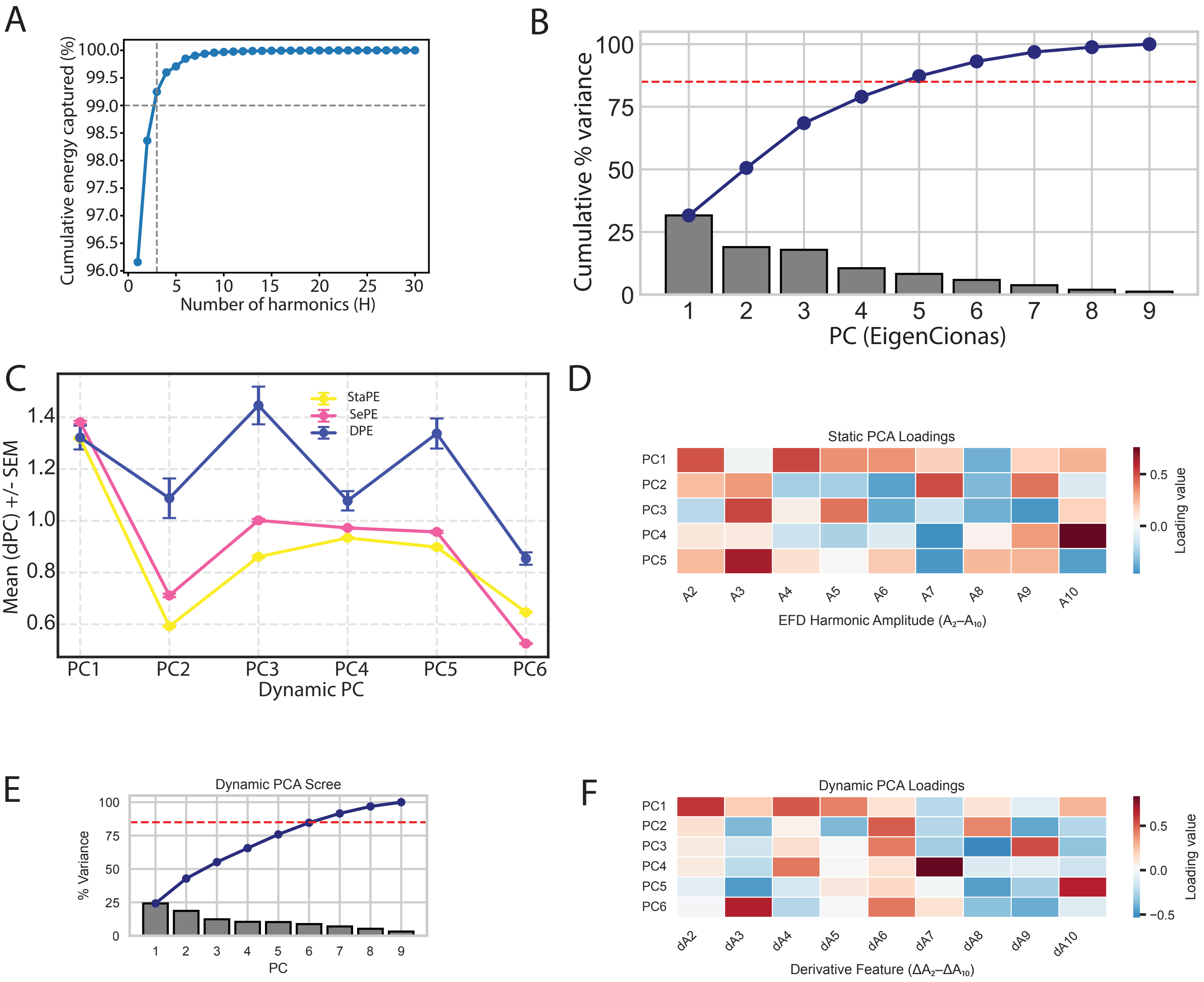
